# How do metal ions direct ribozyme folding?

**DOI:** 10.1101/037895

**Authors:** Natalia A. Denesyuk, D. Thirumalai

**Affiliations:** Institute for Physical Science and Technology, University of Maryland, College Park, Maryland, 20742; Department of Chemistry and Biochemistry and Biophysics Program, University of Maryland, College Park, Maryland 20742

## Abstract

Ribozymes, which carry out phosphoryl transfer reactions, often require Mg^2+^ ions for catalytic activity. The correct folding of the active site and ribozyme tertiary structure is also regulated by metal ions in a manner which is not fully understood. Here, we employ coarse-grained molecular simulations to show that individual structural elements of the group I ribozyme from the bacterium *Azoarcus* form spontaneously in the unfolded ribozyme even at very low Mg^2+^ concentrations, and are transiently stabilized by coordination of Mg2+ ions to specific nucleotides. However, competition for scarce Mg^2+^ and topological constraints arising from chain connectivity prevent complete folding of the ribozyme. A much higher Mg^2+^ concentration is required for complete folding of the ribozyme and stabilization of the active site. When Mg^2+^ is replaced by Ca2+ the ribozyme folds but the active site remains unstable. Our results suggest that group I ribozymes utilize the same interactions with specific metal ligands for both structural stability and chemical activity.

---

Since the remarkable discovery that RNA molecules can function as enzymes^1,2^ an ever increasing repertoire of cellular functions has been associated with these versatile molecules^3^. Execution of these diverse functions, which include control of gene expression and protein synthesis, often requires RNA enzymes (ribozymes) to fold to a compact, functionally competent structure with catalytic metal ions bound at the active site. For example, self-splicing of group I introns is catalyzed by Mg^2+^ ions which coordinate directly to the chemically active RNA groups^4–8^. The close relationship between site-specific Mg^2+^ binding and catalytic activity implies that precise folding of the ribozyme structure is of critical importance. However, folding of the highly negatively charged ribozymes is itself mediated by metal ions^9–12^ using mechanisms that have yet to be fully elucidated^13–20^. In other words, a molecular description of how metal ions facilitate the navigation of the rugged energy landscapes of ribozymes is lacking. Here, we address the problem of ion driven ribozyme folding in computer simulation of the group I intron from the purple bacterium *Azoarcus*^21,22^.

The high-resolution structure of the *Azoarcus* intron is known in complex with two exons in the conformation preceding the second splicing step^7,22,23^ (state pre-2S, Fig. 1). The tertiary structure of the *Azoarcus* intron in the pre-2S state closely resembles the structure of the group I intron from the ciliate *Tetrahymena* in the enzymatic form^24,25^, in which the exons and intron’s internal guide sequence are absent. In our work we have modeled the enzymatic form of the *Azoarcus* intron, assuming that its native conformation is the pre-2S conformation shown in Fig. 1. The crystal structure of the intron shows Mg^2+^ ions located in the regions with high concentration of negatively charged phosphate groups^22,23^. The Mg^2+^ ions in the intron core are either proximal to or directly bound to phosphates that were identified by Tb^3+^ cleavage experiments as candidates for specific interactions with divalent metal ions^12^. Two Mg^2+^ ions in the active site coordinate the reactive phosphate (reactive phosphoryl group), and it has been proposed that they are involved in catalysis^7^. Experiments indicate that high (1 M) concentrations of monovalent ions or submillimolar concentrations of Mg^2+^ ions can cause group I ribozymes to fold into a conformation which is nearly identical to the native state^12,26^. However, the ribozyme catalytic activity requires Mg^2+^ concentration above 1 mM^14^. Based on these findings it was suggested that the unfolded *Azoarcus* ribozyme assembles into inactive compact intermediate, which undergoes subsequent reorganization into the native conformation^14,27^ due to specific binding of Mg^2+^ ions in the ribozyme core. Both the intermediate and native conformations were shown to be stabilized by native tertiary interactions. However, neither the precise structural differences between these conformations nor the role of metal ions in their assembly is understood. Here we use coarse-grained computer simulations to determine the structural properties of the inactive intermediate and, for the first time, to make explicit the relationship between Mg^2+^ coordination and the folding of RNA into an active conformation.

**Figure 1:**
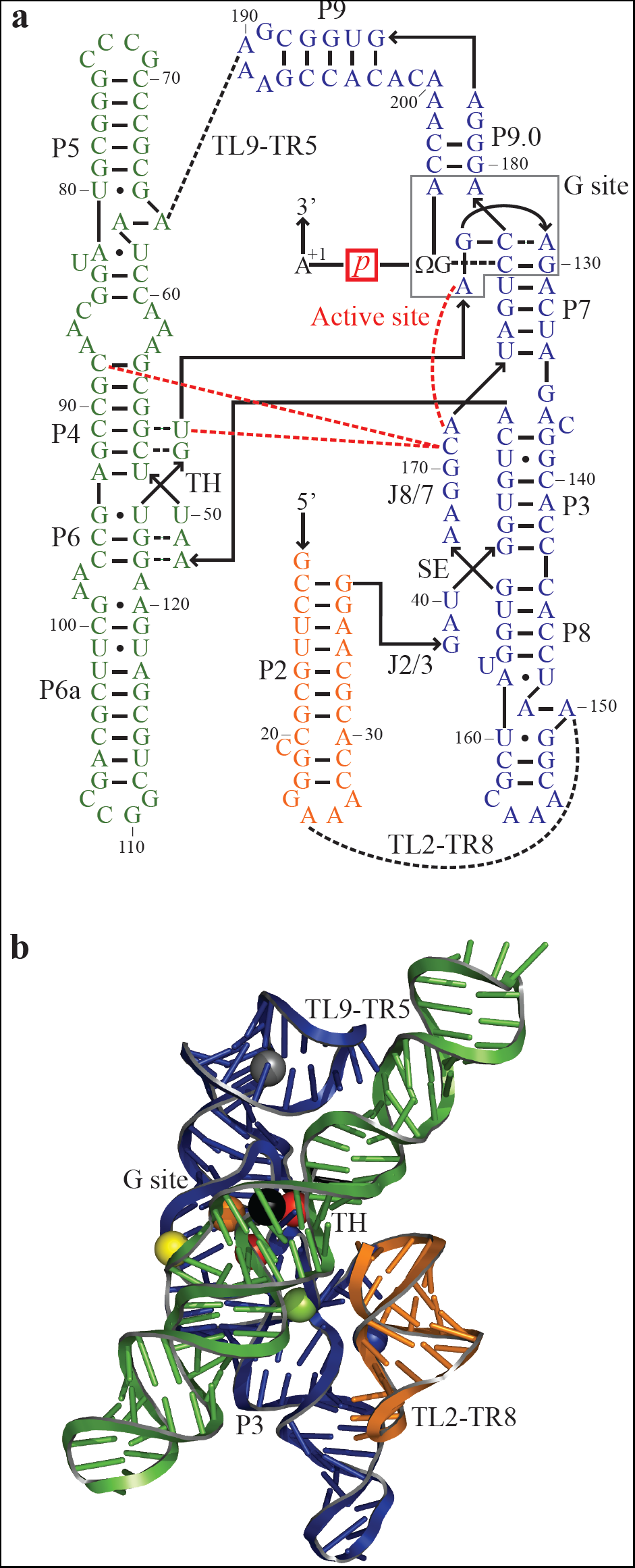
Secondary and tertiary structure of the *Azoarcus* group I intron studied in this work. **a,** Secondary structure, with the intron backbone in solid lines and arrowheads pointing in the 5’ to 3’ direction. The principal secondary structure domains, interdomain junctions, and tertiary interaction motifs are labeled. The gray box encapsulates the nucleotides associated with the G site. Red dashed lines indicate the interdomain interactions stabilizing the active site, which is centered around the reactive phosphate (red box). Other tertiary interactions are shown in black dashed lines. **b,** Three-dimensional crystal structure of the intron^22^ (PDB code 1U6B), using the same colour scheme as in **a.** Spheres indicate the positions of Mg^2+^ ions resolved in the crystal structure that have been shown here to stabilize different tertiary motifs: the G site (orange), TH (black), TL2-TR8 (blue), TL9-TR5 (gray), active site (red), and P3-P7 stacking (yellow). The green sphere shows the predicted position for a Mg^2+^ ion which stabilizes the stack exchange junction SE (not resolved in the crystal structure).

To date an accurate and computationally efficient general simulation technique for studying RNA folding has not been developed. All-atom simulations of RNA in water, originally conceived in the context of protein folding and dynamics, would potentially provide us with the most detailed information on the folding process. However, large uncertainties in atomistic force fields and the difficulty in obtaining adequate conformational sampling have impeded general application of all-atom simulations in the folding studies. Recently it has become possible to generate folding trajectories of a few relatively small proteins in atomistic detail^28–30^. But for RNA, the limitations of all-atom approaches are more formidable, because the tertiary structure of even small ribozymes is known to take from milliseconds to seconds to form. Such long simulation times are not currently possible in all-atom simulations, even using the most advanced technology available. The force fields themselves are known to be inaccurate for the thermodynamics of basic RNA structure formation, such as base stacking^31^. Furthermore, since RNA folding is driven by metal ions, it is essential that experimental ionic conditions be employed in computational studies. In all-atom simulations the ions necessary for the folding of the structure must be contained in a very small simulation box, which implies ion concentrations that far exceed physiological concentrations.

In order to solve the problem of how ions drive ribozyme folding we have developed our own force field for RNA based on a coarse-grained model, in which each nucleotide is replaced by three interaction sites, representing a phosphate, a sugar and a base^32–34^. Our model is one of a class of Gō-like^35^ models for RNA which employ a simplified description of RNA energetics in implicit solvent^32,36,37^. The common simplification used in all G ō-like models is that intramolecular attractive interactions are defined only between the residues that appear to be in contact in the native structure of the RNA molecule. This definition ensures that the native structure of any molecule is the minimum energy structure. By contrast, in all-atom force fields generic attractive potentials are applied to all interatomic pairs and the molecule is not guaranteed to fold into its native structure. The basic drawback of G ō-like models is that they cannot capture any partially folded intermediate states stabilized by non-native interactions. To improve on this approximation, we have gone one step beyond standard G ō-like models and included non-native secondary structure interactions in our RNA model. In particular, we model all base stacking interactions between consecutive nucleotides, as well as hydrogen bond interactions between any bases G (guanine) and C (cytosine), A (adenine) and U (uracil), or G and U. Hydrogen bond and stacking interactions which stabilize the tertiary structure are defined only for the interactions present in the native structure, following the general strategy of G ō-like models. Because secondary structure interactions are substantially stronger than tertiary interactions, we expect that any long-lived misfolded states will be primarily stabilized by non-native secondary structure interactions, and that non-native tertiary interactions would not play a significant role in determining the thermodynamics of ribozyme formation.

Although G ō-like models of RNA have been successful in a variety of applications^32,36–42^, they are typically constructed with a reduced number of energetic parameters, and hence are applicable to a limited range of ion concentrations and temperature. In sharp contrast, the force field used in this study (see Methods) is able to reproduce the experimental thermodynamic and structural data for several different RNA molecules under a relatively wide range of solution conditions. Direct comparisons of the force field predictions with the measured data are presented in Supplementary Methods. The success of the current model in achieving quantitative agreement with experiment is due to a combination of the careful treatment of RNA interactions and explicit inclusion of all ions, which are modeled as spheres characterized by an appropiate charge and radius. This simple description of ions proves to be sufficiently accurate for the aims of the current study, suggesting that the folding of the *Azoarcus* ribozyme is controlled largely by the ion charge density.

## Results and Discussion

### Local and global folding of the *Azoarcus* ribozyme

We report the results of coarse-grained simulations of Mg^2+^-driven folding of the *Azoarcus* ribozyme. The generated equilibrium trajectories are sufficiently long so that we can observe multiple unfolding/refolding of individual tertiary elements in the RNA and track the uptake of Mg^2+^ ions by each element. We focus on the folding of the six principal elements of the ribozyme tertiary structure that undergo distinct folding transitions (Fig. 1): (1) the stack exchange junction, SE, which anchors the native pseudoknot P3, (2) the central triple helix TH, (3-4) two peripheral tetraloop-tetraloop receptor interactions, TL2-TR8 and TL9-TR5, (5) the G site, comprising the G-binding pocket and bound reactive nucleotide ΩG206, and (6) the active site, which is formed by interactions of loop J8/7 with the G site, TH and P4. We could detect stable formation of the G-binding pocket only upon binding of ΩG206 in the pocket, in support of the earlier kinetic studies of G binding^43^, and so we do not regard these as distinct events. Similarly, the junction J2/3 forms concomitantly with TL2-TR8, and hence is not considered to be an independent tertiary motif. The folding transitions of different tertiary motifs proceed in the order illustrated in Supplementary Fig. S1 and contribute to the global compaction of the ribozyme, which appears as a single transition in the radius of gyration *R*_g_ with increasing Mg^2+^ concentration (Fig. 2a). The midpoint, *c*_m_, of the *R*_g_ transition as a function of Mg^2+^ concentration is not sensitive to K^+^ concentration, consistent with the idea that the transition is driven by Mg^2+^ ions. Our results confirm that, in the absence of Mg^2+^, increasing KCl from 12 to 50 mM is sufficient to induce significant reduction in *R*_g_ (Fig. 2a). However, the elements of the tertiary structure do not form in 50 mM KCl without Mg^2+^, except for 17% occurrence of the folded G site (Fig. 2b and Supplementary Fig. S2). In addition to unstable tertiary structure, we find that the helix P3 is unpaired in low Mg^2+^ and its stability curve closely follows that of tertiary motif SE with increasing Mg^2+^ concentration. This and further results for the correlations between secondary and tertiary structure formation can be found in Supplementary Discussion.

**Figure 2:**
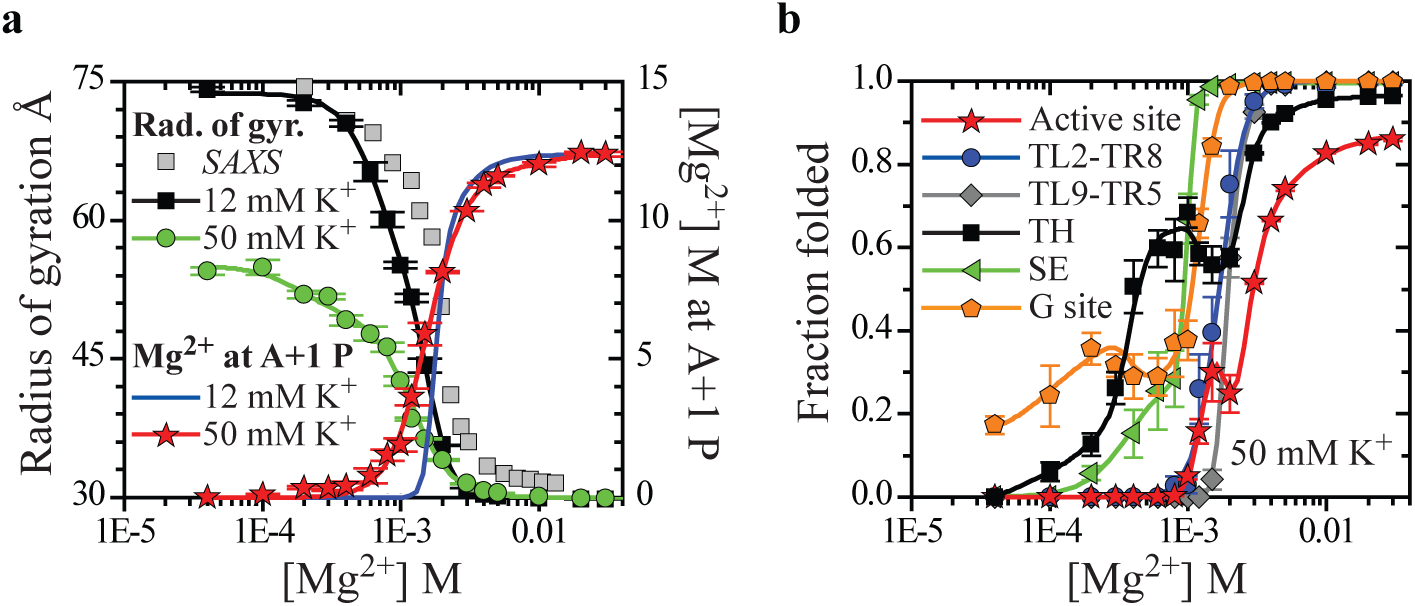
Mg^2+^ promotes folding and function of the *Azoarcus* ribozyme. **a,** The decrease of the radius of gyration with increasing Mg^2+^ concentration indicates global folding of the ribozyme (left axis, black squares — 12 mM K^+^, circles — 50 mM K^+^). Gray squares: radius of gyration obtained in SAXS experiments in 20 mM Tris-HCl buffer^14^ with ionic strength approximately equivalent to 12 mM K^+^. The Mg^2+^ concentrations in the experimental data have been rescaled to compensate for different RNA concentrations used in the simulation and SAXS experiments^45,48^ (details in Supplementary Discussion). The increase of local molar concentration of Mg^2+^ at the surface of the reactive phosphate (right axis, red and blue) indicates local folding of the active site of the functional ribozyme. Around 2 mM Mg^2+^ the ribozyme populates a compact but non-functional state, *I*_c_ (see Supplementary Fig. S9). **b,** Ribozyme tertiary structure folds in two distinct phases. First phase folding, below 1 mM Mg^2+^: G site (orange), stack exhange junction SE (green), base triple G53-C91-U126 in the TH (60% folded, black). Second phase folding, above 1 mM Mg^2+^: G53-C91-U126 (100% folded, black), peripheral contacts TL2-TR8 (blue) and TL9-TR5 (gray), contacts between J8/7 and P4 at the active site (red). Folding curves for additional elements of the TH and ribozyme core are plotted in Supplementary Fig. S5. The origin of the nonmonotonic Mg^2+^ dependence for the active site, G site and TH is elucidated in Supplementary Fig. S6. Error bars show standard deviations obtained from the bootstrap analysis of the primary data set in which each data point represents the mean from a 7.5 sampling interval.

### Mg^2+^ coordination of the folded ribozyme

The spatial distributions of Mg^2+^ ions at different stages in the ribozyme assembly contain pronounced peaks at RNA sites characterized by high affinity for Mg^2+^ (Fig. 3). These Mg^2+^ concentration profiles are fingerprints, which identify site specific ion-RNA interactions that direct the folding of tertiary structure. In 30 mM Mg^2+^, the highest concentration considered, the *Azoarcus* ribozyme is fully folded (Fig. 2a, b). The majority of high-affinity sites in the Mg^2+^ fingerprint at 30 mM (Fig. 3a) are consistent with the positions of Mg^2+^ ions resolved in the crystal structure of the intron^22,23^. At the lowest Mg^2+^ concentration at which the ribozyme is folded (4 mM, Fig. 2a, b), the local Mg^2+^ ion concentration at the high-affinity sites is the same as in 30 mM, whereas it decreases noticeably elsewhere (Fig. 3a). This indicates that the ribozyme tertiary structure is sustained by localized Mg^2+^ ions.

**Figure 3:**
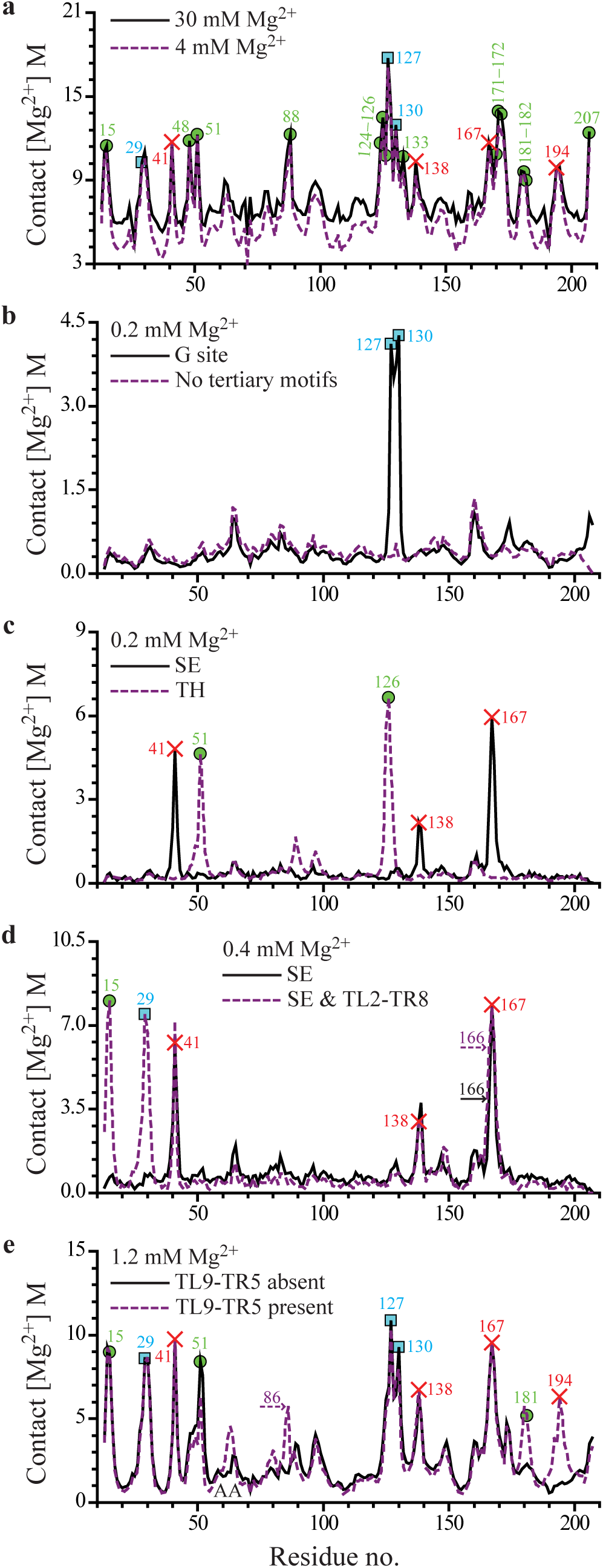
Mg^2+^ fingerprints in 50 mM KCl. Contact Mg^2+^ concentration, j-axis, is the local molar concentration of Mg^2+^ at the surface of phosphate groups (quantitative definition in Supplementary Fig. S10). Symbols mark nucleotides that are either in direct contact with Mg^2+^(circles) or linked to an ion via a water molecule (squares) in the crystal structure^22^. Crosses mark Mg^2+^ coordination sites identified here which do not have an analogue in the crystal structure. The first nucleotide in the exon sequence, A+1, is labeled 207. **a,** Mg^2+^ fingerprints of the fully folded ribozyme in 4 mM (dashed line) and 30 mM Mg^2+^ (solid line) indicate sites with high affinity for Mg^2+^. In panels **b-e,** these high affinity sites are traced to individual tertiary motifs. **b,** Solid line: Mg^2+^ fingerprint of the G site in class 1 RNA conformations at 0.2 mM. In the absence of any folded tertiary motif (class 6 conformations), the ribozyme does not have sites with high affinity for Mg^2+^ (dashed line). **c,** Mg^2+^ fingerprints of single motifs SE (class 3, solid line) and TH (class 2, dashed line) at 0.2 mM. Six conformational classes at 0.2 mM are introduced in the main text. **d,** Mg^2+^ fingerprint of simultaneously folded SE and TL2-TR8 (dashed line) vs. fingerprint of SE alone (solid line) at 0.4 mM. Comparison of peaks establishes a Mg^2+^ coordination pattern for TL2-TR8. Horizontal arrows mark the increase in the coordination level of G166 due to the folding of TL2-TR8. **e,** Mg^2+^ fingerprints for the ribozyme conformations with folded G site, SE, TH, TL2-TR8 and either unfolded (solid line) or folded (dashed line) TL9-TR5 at 1.2 mM. In addition to numbered peaks, the formation of TL5-TR9 promotes the accumulation of Mg^2+^ near stacking platform A63-A64 (AA).

### Mg^2+^ coordination of the unfolded ribozyme

The Mg^2+^ fingerprints in Fig. 3 reveal that the distinct subsets of the peaks, associated with the formation of each of the tertiary motifs, emerge as Mg^2+^ concentration is gradually increased. The presence of very few Mg^2+^ ions per RNA is sufficient to trigger folding of the G site, SE and TH in the unfolded ribozyme. In submillimolar Mg^2+^ and 50 mM KCl these tertiary motifs form intermittently, as illustrated by the equilibrium trajectories in Supplementary Fig. S3. At 0.2 mM Mg^2+^ the ensemble of RNA conformations partitions into six structural classes characterized by the formation of the G site, SE and TH: (1) in 33% of conformations only the G site is formed, (2) in 12% of conformations only TH, (3) in 2.2% only SE, (4) in 2.8% the G site and SE, (5) in 0.7% SE and TH, (6) and in the remainder of the conformations none of the three motifs are formed. As we will discuss below, the complete folding of the G site and TH is mutually exclusive in submillimolar Mg^2+^ due to a phenomenon which we call folding frustration. Folding frustration occurs if topological restrictions arising from chain connectivity prevent the free energies of all interaction sites from being simultaneously minimized.

The Mg^2+^ fingerprint characteristic of class 1 RNA conformations with the folded G site has sharp maxima at residues A127 and G130 (Fig. 3b). A similar Mg^2+^-RNA interaction pattern is also observed in the crystal structure of the folded intron^22,23^, where a single Mg^2+^ ion is coordinated via a water molecule to phosphates 127 and 130, which are at distances 5.7 Å and 6.6 Å from the ion, respectively. The formation of the stack exchange junction SE in the absence of other motifs (class 3 conformations) is accompanied by the accumulation of Mg^2+^ ions in a cavity lined by phosphates 41, 138 and 167-169 (Fig. 3c and Supplementary Fig. S4). There are no resolved Mg^2+^ ions in the cavity in the crystal structure of the folded intron, possibly due to the diffuse nature of the local ion distribution. The Mg^2+^ fingerprint of the triple helix TH in class 2 conformations has substantial peaks around nucleotides 51 and 126 (Fig. 3c). We associate these peaks with two Mg^2+^ ions resolved in the crystal structure, localized at distances 3.8 Å and 2.6 Å from phosphates 51 and 126, respectively. Analysis of individual folding events (Supplementary Fig. S3) indicates that the appearances of the unique Mg^2+^ fingerprints in Fig. 3b, c are precisely correlated in time with the formation of the corresponding tertiary motifs. When none of the tertiary structure is formed (class 6 conformations), the ribozyme does not have sites with high affinity for Mg^2+^ (Fig. 3b). This is consistent with our finding that coordination of Mg^2+^ ions to transiently formed tertiary motifs initiates folding of the ribozyme structure.

Of the three tertiary motifs, the triple helix TH shows the strongest affinity for Mg^2+^ (Fig. 3c) and, consequently, the largest increase in stability in submillimolar Mg^2+^ (Fig. 2b). Interestingly, in the absence of Mg^2+^, the motif SE is most stable, indicating that it is not the intrinsic stability of a tertiary structure which determines its ability to capture Mg^2+^ ions. This ability depends primarily on the details of the electrostatic potential of the phosphate-lined recruiting pockets associated with structurally diverse tertiary motifs.

### Mg^2+^ fingerprints of the peripheral motifs

In some cases, folding of a tertiary structure is a prerequisite for subsequent folding of another motif. For example, the tetraloop-tetraloop receptor interaction between domains P2 and P8 (TL2-TR8) can be detected only when the stack exchange junction SE is correctly folded. Figure 3d illustrates the Mg^2+^ fingerprint corresponding to simultaneous presence of folded SE and TL2-TR8 in 0.4 mM Mg^2+^. Comparison with the Mg^2+^ fingerprint of SE establishes Mg^2+^ binding region associated with TL2-TR8 itself (Figs 3d and 4a). The predicted region, occupied by a Mg^2+^ ion in the crystal structure of the intron (Fig. 4a), is not proximal to the P2 tetraloop, indicating that Mg^2+^ ions stabilize tertiary interactions indirectly.

**Figure 4:**
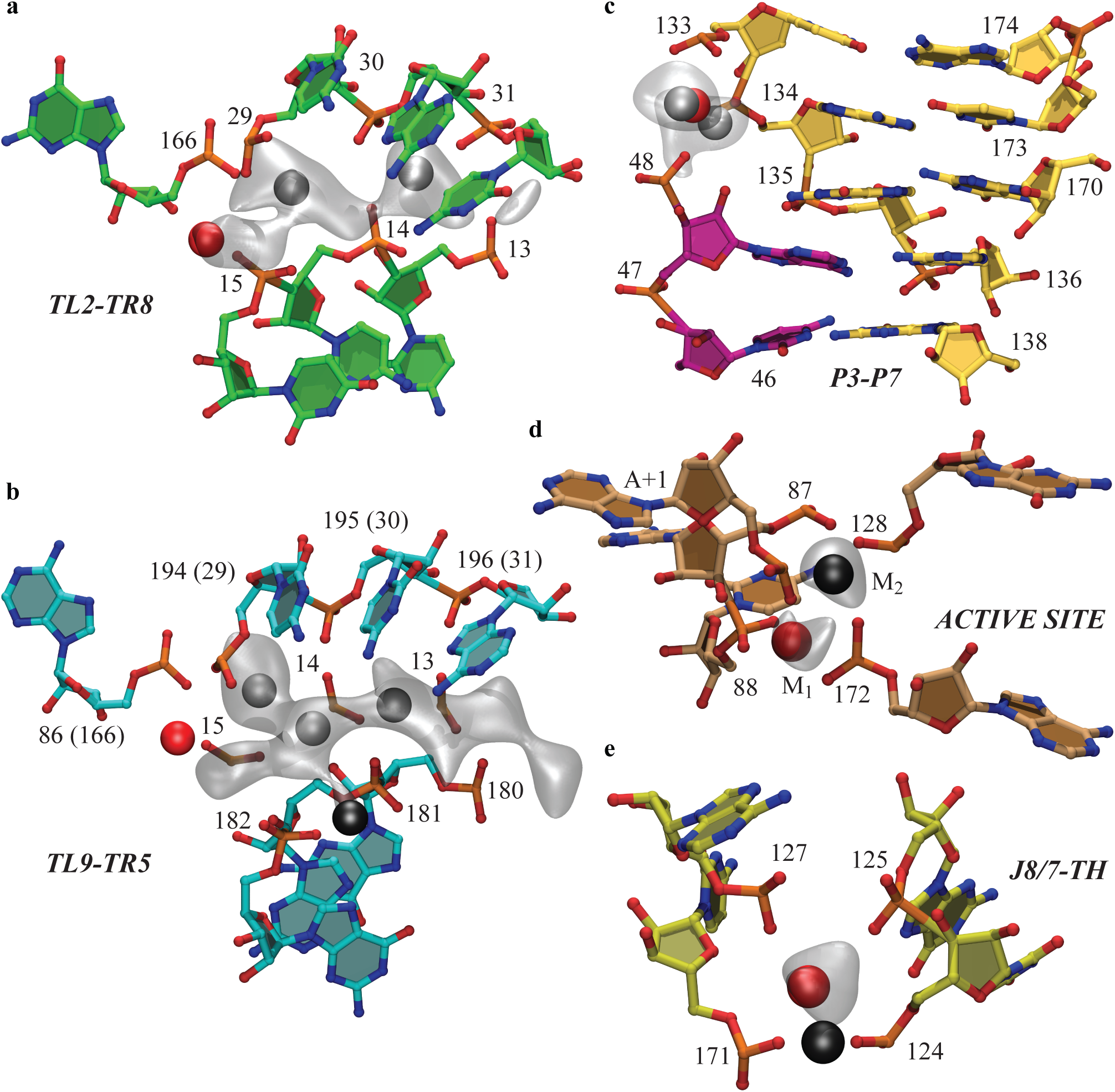
Site specific Mg^2+^-RNA interactions stabilize principal tertiary structure motifs. Gray clouds show regions of local Mg^2+^ concentration exceeding 50% of its maximum value for each motif. For some motifs local Mg^2+^ concentration has several maxima of equivalent height (gray spheres). **a,** A fragment from motif TL2-TR8 characterized by high affinity for Mg^2+^. High affinity residues are distinct from the residues that establish direct contact between domains P2 and P8. **b,** Analogous fragment from motif TL5-TR9. TL2-TR8 was aligned with TL5-TR9 based on comparison of G166 with A86 and C29-C31 with G194-A196 and the resulting phosphate positions 13-15 were superimposed on the TL5-TR9 fragment. The phosphate groups in TL5-TR9 are further apart (compare G180-G182 to superimposed phosphates 13-15) and the Mg^2+^ cloud is more diffuse than for TL2-TR8. In the crystal structure^22^ one Mg^2+^ ion is bound to TL2-TR8 (red spheres in **a** and **b)** and one to TL5-TR9 (black sphere in **b). c,** Coaxial stacking of domains P3 and P7 is stabilized by Mg^2+^ localized between residues 48 and 133-134. Red sphere is a crystallographic Mg^2+^ ion^22^. **d,** Distribution of Mg^2+^ at the active site has two maxima consistent with the positions of two catalytic Mg^2+^ ions (spheres M1 and M2) in the crystal structure of the active intron^7^. Only M1 is present in the structure of the inactive intron^22^. **e,** Another Mg^2+^ site in the ribozyme core forms simultaneously with the active site. Red and black spheres show alternative positions of a Mg^2+^ ion bound at the site in the crystal structures of the active and inactive introns, respectively.

The Mg^2+^-RNA interaction pattern of TL9-TR5 is similar with that of TL2-TR8. The TL9-TR5 motif unfolds rapidly in submillimolar Mg^2+^, which complicates the determination of its Mg^2+^ fingerprint. Comparison of multi-motif fingerprints in 1.2 mM Mg^2+^ in the absence and presence of TL9-TR5 folding (Fig. 3e) reveals a diffuse Mg^2+^ binding region associated with TL9-TR5 (Fig. 4b). Similarities between Mg^2+^ coordination of TL2-TR8 and TL9-TR5 become apparent upon structural alignment of these motifs (Fig. 4a, b). In the crystal structure, a Mg^2+^ ion is found at the periphery of the TL9-TR5 binding region (Fig. 4b), corroborating the diffuse character of the Mg^2+^ ion distribution observed in simulations.

### Final stage of folding

The peaks around phosphates 48, 88, 124-125, 133, 171-172 and +1 (207) in the Mg^2+^ fingerprints in Fig. 3a are not associated with the G site, SE, TH, TL2-TR8 or TL9-TR5, but emerge cooperatively above 1 mM Mg^2+^. The Mg^2+^ ion coordination with phosphates 48 and 133, also found in the crystal structure, stabilizes coaxial stacking of the native pseudoknot P3 and helix P7 (Fig. 4c). The probability for these domains to stack coaxially increases with Mg^2+^ concentration with an approximate midpoint of 1.5 mM (Supplementary Fig. S5), which is noticeably higher than the midpoint for coaxial stacking of P3 and P8 (SE in Fig. 2b). The peaks involving phosphates 88, 124-125, 171-172, 207 are associated with the formation of tertiary contacts in the ribozyme active site. In support of this, the growth of the peak at reactive phosphate 207 with increasing Mg^2+^ concentration (red and blue curves in Fig. 2a) parallels the folding curve of the active site (red curve in Fig. 2b and Supplementary Fig. S2). Analysis of the spatial distribution of Mg^2+^ ions at the active site shows that it is strongly localized around two distinct maxima (Fig. 4d). These maxima are consistent with two Mg^2+^ ions bound at the active site in the crystal structure of the intron, which are essential to the catalytic activity^7^. This demonstrates that the Mg^2+^ ions at the active site serve the dual purpose of structural stabilization and catalysis (Fig. 4d). The active site forms cooperatively with a complementary electronegative pocket, lined by phosphates 124-125, 127 and 171 (Fig. 4e). A Mg^2+^ ion occupying this pocket, observed in simulations as well as in the crystal structure (Fig. 4e), further contributes to active site stabilization. A cooperative link between the two sites has also been confirmed by Tb^3+^ cleavage experiments that pointed to the ability of nucleotide 171 to bind Mg^2+^ and switch the ribozyme from the inactive to the active state^12^.

### Anti-cooperativity and cooperativity of tertiary interactions

Interactions between tertiary motifs cause the stability of some motifs to change nonmonotonically with Mg^2+^concentration (Fig. 2b and Supplementary Fig. S2). One example is interaction between the G site and TH which are linked directly by the RNA chain, resulting in the folding frustration and anti-cooperativity between these motifs. For the G site and TH to be simultaneously folded, residues U126-A127 must adopt an entropically unfavorable extended conformation (Fig. 1a). Consequently, at less than 2 mM Mg^2+^, the stability of the base triple G53-C91-U126 in the TH decreases when the stability of the G site increases, and vice versa (Fig. 2b and Supplementary Figs S2 and S6). The stability of the base triple C52-G92-G125 in the TH is also dependent, to a lesser extent, on the formation of the G site (Supplementary Fig. S5). Only when Mg^2+^concentration exceeds 4 mM its stabilizing effect on the TH overcomes the anti-cooperativity effects and the folding of G53-C91-U126 can proceed to completion (Fig. 2b and Supplementary Fig. S2). We find that, despite strong anti-cooperative correlation between the TH and G site, the folded TH in fact stabilizes other elements of the ribozyme tertiary structure (see Supplementary Discussion).

We have also observed a destabilizing effect of the peripheral motif TL9-TR5 on the interactions of J8/7 with P4 and TH, which causes the stability of the active site to decrease around the folding transition midpoint of TL9-TR5 (Fig. 2b and Supplementary Figs S2, S5 and S6). The folding of the active site and the folding of TL9-TR5 draw together coaxially stacked domains P5-P4-P6a and P7-P3-P8 in mutually inconsistent relative orientations (Supplementary Fig. S7). When neither active site nor TL9-TR5 are folded, the angle between the two domains, *γ*, undergoes large fluctuations with mean close to 70° (Fig. 5a and Supplementary Fig. S2). The formation of the active site alone increases the mean value of *γ* to approximately 90°. In contrast, the folding of TL9-TR5 in the absence of the folded active site results in a relatively narrow distribution of *γ* with mean below 60° (Fig. 5a and Supplementary Fig. S2). As a consequence of this conformational conflict between the active site and TL9-TR5, only one of the two motifs is observed in the majority of ribozyme conformations below 3 mM Mg^2+^. Increasing Mg^2+^ concentration above 4 mM leads to significant population of the native conformation, in which all core and peripheral interdomain contacts are formed and *γ* is narrowly distributed around 65° (Fig. 5a and Supplementary Fig. S1), as compared to 67° in the crystal structure^22^. In the native conformation the compromise value of *γ* is attained through the formation of a kink between helices P9 and P9.0 (Fig. 1). At 30 mM Mg^2+^ the native conformation is populated less than 100% due to lack of complete stability of the active site (Supplementary Fig. S5). It is possible that the conformational mobility in the ribozyme core in the absence of substrate is necessary for efficient substrate binding and catalytic function.

**Figure 5:**
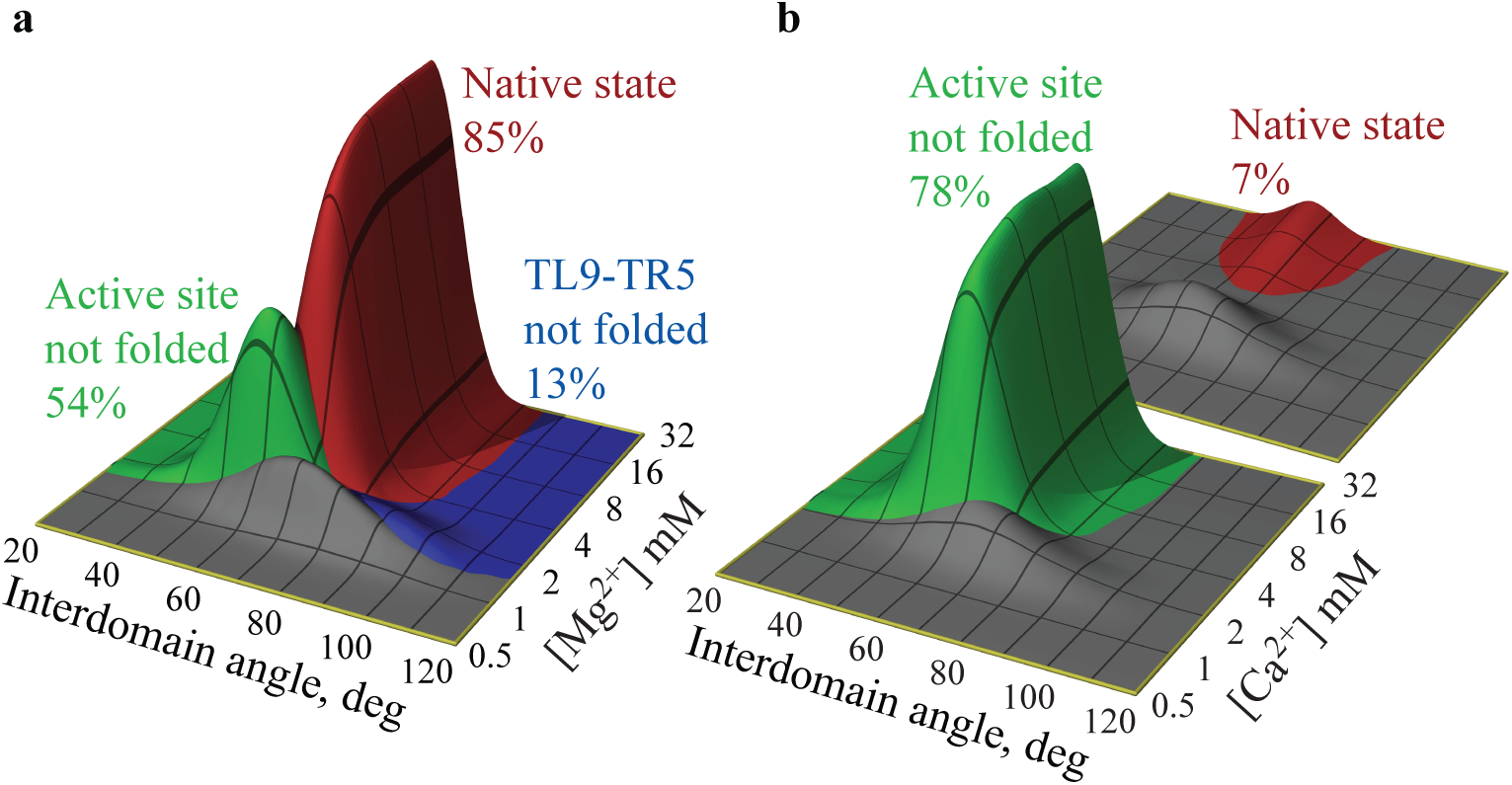
Conformational states of the *Azoarcus* ribozyme at intermediate to high concentrations of divalent ion. **a,** The ribozyme is in one of four states characterized by the presence of rigid domains P5-P4-P6a and P7-P3-P8 interacting at: (1) the active site and peripheral junction TL9-TR5 (native state, red), (2) active site only (blue), (3) TL9-TR5 only (green), (4) neither (gray). The four states are represented by their respective, Mg^2+^-dependent, probability distributions of the angle *γ* formed by domains P5-P4-P6a and P7-P3-P8 (*γ* is defined in Supplementary Fig. S7). The percentages in the panel indicate the maximum occupancy of states 1-3 amid all Mg^2+^ concentrations in 50 mM K^+^ (12 mM K^+^ data are in Supplementary Fig. S2). The maximum occupancy of state 4 is 30%. Together, states 2-4 represent the experimentally observed inactive compact state *I*_c_. **b,** Same as in panel **a** but for the folding in Ca^2+^. The ribozyme active site is unstable in the absence of Mg^2+^, which largely eliminates the native state (shown at rear for clarity of representation) and state 2 (1% occupancy, not shown). In the limit of high Ca^2+^, the ribozyme adopts the conformation analogous to state 3 in panel **a.** The maximum occupancy of state 4 is 19%.

We propose that an ensemble of conformations in which the principal domains P5-P4-P6a and P7-P3-P8 are formed but are not in the native orientation represents the native-like inactive intermediates, *I*_c_, observed experimentally^44,45^ (gray, blue, green in Fig. 5a and Supplementary Fig. S2). Specific details of this conformational ensemble depend on the concentration of the monovalent ion. High concentration of KCl enhances the stability of tetraloop-tetraloop receptor motifs, thus promoting the conformations with folded TL9-TR5 while decreasing the population of conformations with the folded active site (Fig. 5a and Supplementary Fig. S2). It should be possible to distinguish various conformational states and confirm this prediction using FRET experiments in which one pair of fluorescent markers is attached to nucleotides forming a peripheral contact between P5 and P9, and another pair to nucleotides connecting J8/7 and P4 or J8/7 and TH (Fig. 1a). The proposed nature of the native-like intermediates explains why the motif TL9-TR5 is unfolded in one of the four molecules comprising the unit cell in the crystal structure of the *Tetrahymena* ribozyme^25^.

Early studies of the unfolding of group I intron of bacteriophage T4 described the cooperative loss of most of the tertiary interactions upon heating, which appeared as a single twōstate transition^46^. More recent mutation studies of the *Azoarcus* ribozyme folding concluded that the initial population of the native-like intermediates is also guided by a cooperative network of tertiary interactions with the TH helix at its center^14^. Surprisingly, strong anti-cooperativity between the peripheral motif TL9-TR5 and interdomain contacts in the ribozyme core emerges as the folding progresses to the native structure at higher Mg^2+^concentration^14^. These results are in complete accord with our simulations which place partial formation of the TH at the beginning and coexistence of TL9-TR5 and the active site during the final stage of ribozyme folding with increasing Mg^2+^ concentration. We find that it is precisely the folding frustration between the core and peripheral tertiary contacts, characteristic of the native conformation, that leads to the population of native-like states at intermediate Mg^2+^ concentration. The possibility that not all elements of RNA tertiary structure are linked cooperatively was also discussed in the context of kinetic studies of the G binding pocket^43^, which was shown to undergo a distinct folding transition upon binding of guanosine. The results presented here provide support and the much-needed structural underpinning for these insightful experiments.

### Folding in Ca^2+^

To examine the specific requirement of Mg + for the catalytically competent assembly of the *Azoarcus* ribozyme, we carried out additional simulations using Ca^2+^ instead of Mg^2+^. Our results for the tertiary structural equilibria in 50 mM KCl with varying Ca^2+^concentration, summarized in Supplementary Fig. S8, show that the folding transition midpoint in Ca^2+^ is higher than in Mg^2+^ and the ribozyme folded state is less compact. We find that most of the tertiary structure has formed in Ca^2+^ with the exception of the active site and the base triple G53-C91-U126 in the TH (Supplementary Fig. S8). In addition, the probability for P3 and P7 to stack coaxially is less than 100% in 30 mM Ca^2+^ (Supplementary Fig. S8), pointing to the presence of conformations with incompletely formed domain P7-P3-P8. The majority of compact conformations in Ca^2+^ are similar to an intermediate observed for Mg^2+^, in which domains P5-P4-P6a and P7-P3-P8 are completely formed and joined by the TL9-TR5 but not J8/7-P4 or J8/7-TH contacts (green in Fig. 5a, b). The probability of formation of the native conformation, representing a potentially active ribozyme, is low in 30 mM Ca^2+^ (Fig. 5b).

The stark difference in the stability of the active site in Mg^2+^ and Ca^2+^ cannot be attributed to the electrostatic energy alone. Indeed, the energy of Coulomb interaction between a phosphate group and a Ca^2+^ ion at the distance of closest approach is only 25% less than the analogous energy for a Mg^2+^ ion. This leads us to conclude that the most important discriminating factor between Mg^2+^ and Ca^2+^ ions is not the Coulomb energy, but the size exclusion of larger Ca^2+^ ions from the active site. In the crystal structure of the active intron^7^, two Mg^2+^ ions bound at the active site are separated by 3.9 Å (Fig. 4d). This short distance is only slightly larger than the diameter of a single Ca^2+^ ion, indicating that the active site cannot accommodate two Ca^2+^ ions without significant steric frustration between them. Similarly, in Fig. 4e the radius of gyration of the electronegative binding pocket adjacent to the active site is 3.8 Å, which just equals the sum of the phosphate and Ca^2+^ radii. A Ca^2+^ ion cannot effectively bind in such a tight configuration without disturbing the binding pocket itself and neighboring RNA structure.

## Conclusions

Our study establishes that site specific interactions between Mg^2+^ ions and individual tertiary motifs occur even in the unfolded ribozyme. Due to this extraordinary specificity, as few as two Mg^2+^ ions per RNA (0.1 mM Mg^2+^ in Fig. 2b) can serve to nucleate transient folding of key tertiary motifs. At such low Mg^2+^ concentrations the folding of the tertiary motifs is mutually exclusive since the ions must be released before another motif can fold. With increasing Mg^2+^ concentration, multiple motifs fold in parallel in accordance with their affinity for Mg^2+^ ions and subject to the topological constraints of the RNA. A complex order of equilibrium assembly arises from these interactions, with the stability of some tertiary structure changing non-monotonically with Mg^2+^ concentration (Fig. 2b). Although the principal helical domains in the *Azoarcus* ribozyme can also fold in Ca^2+^, their correct relative orientation and the organization of the active site require Mg^2+^ ions, which have a much higher charge density. Our results demonstrate that the Mg^2+^ coordination pattern necessary for catalytic activity also provides the basis for structural stability of the active site (Fig. 4d). This conclusion is further supported by our all-atom simulations described in Supplementary Discussion, as well as by experiment^12^. Such harmony between chemical and structural requirements reduces the possibility that the active site is folded and occupied by catalytically inactive ions which must be displaced before the splicing reaction can proceed. Only the Mg^2+^-coordinated active site is ordered, poising the ribozyme for substrate recognition and catalytic activity, thus effectively speeding up the rate of reaction.

The folding mechanism we have discovered — in which folding of individual structural elements results in formation of phosphate-lined binding pockets that recruit stabilizing Mg^2+^ ions, even at Mg^2+^ concentrations for which global folding is frustrated — is likely to be quite general. The interactions between Mg^2+^ ions and the RNA are determined by the structural properties of these binding pockets, rather than by a specific RNA sequence. Furthermore, homology between the structural elements in the ribozyme studied here and many other functional RNA molecules suggests that the relationship between Mg^2+^ binding and folding elucidated here should hold in other ribozymes.

## Methods

The force field used in this study is an extension of our earlier model^34^. It takes into account bond length and valence angle constraints, secondary and tertiary structure hydrogen bonding, secondary and tertiary structure base stacking, excluded volume repulsions and electrostatic interactions. Below we summarize the new elements in the interaction potentials that were introduced to address the problem of folding of a large ribozyme.

The potentials for bond lengths and valence angles are carried over from the original model^34^. Both models incorporate the hydrogen bonds present in the crystal structure of the RNA molecule, which are determined by submitting the structure to the WHAT IF server at http://swift.cmbi.ru.nl. In the current model other non-native canonical base pairs can form between any A and U, G and C, and G and U separated by at least four nucleotides along the chain. The potential for hydrogen bonds, which was 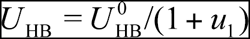 in the original model, is 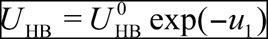 in the new model, where the common function *u*_1_ is a combination of harmonic potentials chosen to bias the structure to ideal A-form helix for canonical bonds or to the crystal structure for non-canonical bonds^34^. The rapidly decaying exponential form of the revised *U*_HB_ describes the short range of hydrogen bonds more accurately, which is important when there is a large number of non-native interactions. 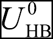 is an adjustable parameter.

Stacking interactions between two consecutive nucleotides are modeled using the same potential in both models, 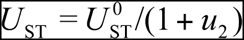, where *u*_2_ is a sum of harmonic terms that bias the stack structure to A-form helix^34^. The parameters 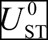 take different values for sixteen (16) different nucleotide dimers, r(XpY), where X, Y represents A, C, G, or U. The 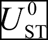 are temperature-dependent, 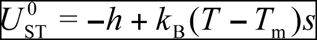, where *h* and *s* are tuned for each r(XpY) individually so as to yield the melting temperatures (*T*_m_) and entropies of r(XpY) stacking obtained from experiment, as detailed previously^34^. The parameters *h* (but not *s*) which result from this learning procedure are functions of a single free energy correction ΔG_0_^34^. ΔG_0_ is the second adjustable parameter.

Tertiary stacking interactions between nonconsecutive nucleotides were not included in the original model. We have identified twenty seven (27) stacks between nonconsecutive nucleotides in the crystal structure of the *Azoarcus* intron^22^ (Supplementary Table S1). These stacks are described here using the interaction potential 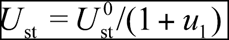, where *u*_1_ is the same as for hydrogen bonds. Note that native tertiary stacks are modeled similarly to native hydrogen bonds in the original model, because there are no fundamental differences in the geometry of base pairing and base stacking in a coarse-grained representation of RNA. *U*_st_ is the third adjustable parameter.

All electrostatic interactions are modeled using Coulomb potential, divided by the temperature dependent dielectric constant of water^47^. Solvent molecules are not explicitly included in simulation. The charges for RNA sites and ions are given in Supplementary Table S2. In the original force field the ions were modeled implicitly^34^.

Excluded volume repulsion between sites *i* and *j* (RNA sites or ions) separated by distance *r* (Å) is described by the modified Lennard-Jones potential,

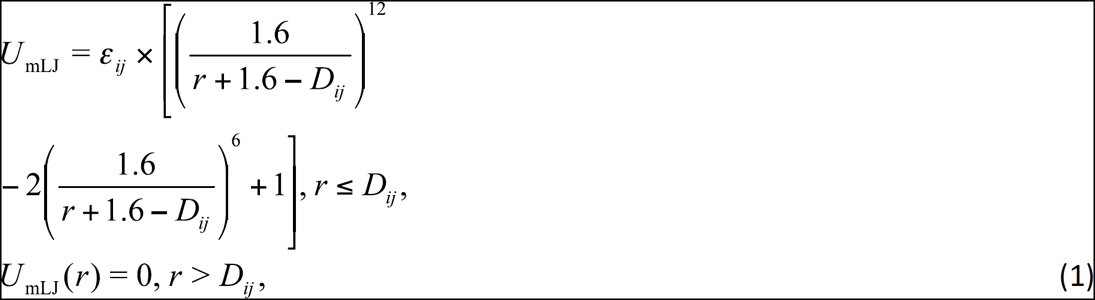

where *D_ij_* = *R_i_* + *R_j_* and 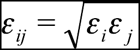. *R_i_ ε_i_* and for ions and RNA are listed in Supplementary
Table S2. *U*_mLJ_ models hard repulsions that decay on the short length scale of 1.6 Å. The use of *U*_mLJ_ simplifies parameterization of the model, because the short range of repulsions makes our quantitative results insensitive to the specific values of *ε_i_*. The distance 1.6 Å ensures that *U*_mLJ_ becomes a standard Lennard-Jones potential for a pair of smallest particles — two Mg^2+^ ions. *R_i_* for RNA have been adapted from the original model^34^. In Supplementary Methods we provide justification for our choice of *R_i_* for divalent ions, and also demonstrate that RNA thermodynamics is relatively insensitive to *R_i_* for K^+^.

Using comparison of simulation and experimental melting curves for an RNA hairpin and pseudoknot we have set 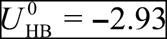 kcal/mol, ΔG_0_ = 0.85 kcal/mol and 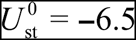 kcal/mol (Supplementary Methods). In the original model ΔG_0_ = 0.6 kcal/mol^34^, which results in small differences in *h* between the two models (*s* are the same). *h* and *s* for sixteen dimers are listed in Supplementary Table S3.

The proposed force field may be applied to other RNA molecules provided the structure of RNA is available to determine a network and geometric parameters of non-canonical hydrogen bonds and nonconsecutive stacks, as was done in the simulations reported here. Further details of these simulations can be found in Supplementary Methods.

## Author contributions

N.A.D. and D.T. conceived and designed the project, analyzed the simulation data and cōwrote the paper. N.A.D. performed the simulations.

## Acknowledgements

This work was supported by a grant from the National Science Foundation (CHE 13-61946). Correspondence and requests for materials should be addressed to D.T.

